# CO_2_ signalling mediates neurovascular coupling in the cerebral cortex

**DOI:** 10.1101/2020.12.31.424942

**Authors:** Patrick S Hosford, Jack A Wells, Iván Ruminot, Isabel N Christie, Shefeeq M Theparambil, Anna Hadjihambi, L Felipe Barros, Mark F Lythgoe, Alexander V Gourine

## Abstract

The mechanisms of neurovascular coupling remain incompletely understood. Here we show in experimental animal models that the neuronal activity-dependent increases in local cerebral blood flow in the somatosensory cortex are abolished by saturation of brain CO_2_-sensitive vasodilatory mechanism or disruption of brain HCO_3_^-^/CO_2_ transport, independently of baseline cerebral perfusion and brain tissue pH. These results suggest that increases in metabolic CO_2_ production and CO_2_ signalling play an important role in the development of neurovascular coupling response.

## Introduction

Neurovascular coupling is a fundamental, evolutionarily-conserved signalling mechanism responsible for dilation of cerebral blood vessels and increase in local cerebral blood flow (CBF) in response to heightened neuronal activity. The main purpose of the neurovascular response is thought to be to maintain an uninterrupted delivery of oxygen and glucose to support changing neuronal metabolic needs. The methods of functional magnetic resonance imaging (fMRI) record these changes in local CBF and use them as a proxy of neuronal activity to study the human brain. Most recent evidence suggests that long-term impairment of neurovascular coupling may precipitate age-related neuronal damage, contribute to cognitive decline and the development of neurodegenerative disease^1,2^. Therefore, a full understanding of the signalling mechanisms between neurons and the cerebral vasculature may prove to be critically important for the development of future treatments of these conditions, as well as for our understanding and interpretation of human brain imaging data.

However, there are still controversies surrounding the functional significance and the mechanisms underlying the neurovascular coupling response^3,4^. In accord with the currently accepted view, neurovascular response is driven by neuronal activity-induced changes in the brain neurochemical milieu (neurotransmitter spillover, accumulation of K^+^) leading to activation of intermediate cell types, including interneurons and astrocytes, which in turn signal to vascular smooth muscle cells and pericytes^5^. A recent meta-analysis of published data reporting the effects of pharmacological or genetic blockade of all hypothesized signalling pathways indicated that such feed-forward mechanisms may account for up to ~60% of the neurovascular response^3^. The analysis pointed to the existence of an as yet unidentified signalling mechanism(s) responsible for a significant proportion (at least one third) of the response^3^.

In air-breathing animals with a high metabolic rate living in conditions of ample oxygen supply, the effective removal of metabolically produced CO_2_ is critical to maintain homeostasis. The human brain generates ~20% of total body CO_2_ production (3.3 moles or ~75 litres per day) and this CO_2_ can only be removed from the brain by the cerebral circulation. All membrane, molecular and biochemical processes involved in synaptic transmission are affected by changes in pH, therefore, uncontrolled fluctuations in brain tissue CO_2_/pH are detrimental to neuronal function. CO_2_ production increases in parallel with the neuronal activity and energy use^6^ and has an extremely potent dilatory effect on the brain vasculature^7^, which (in contrast to systemic vessels) is uniquely sensitive to CO_2_^8^. Conceivably, the need for effective removal of surplus CO_2_ generated during increased brain activity and brain pH regulation was an important driving force behind the evolutionary development of the neurovascular coupling mechanism. If so, CO_2_ may act as a signalling molecule between active neurons and brain vessels. Yet, rather surprisingly, the role of locally produced CO_2_ in the neurovascular coupling response has never been experimentally addressed.

## Results and Discussion

To test this hypothesis, methods of specific blockade of CO_2_ transport or actions in the brain *in vivo* are required. However, as the exact mechanism of the CO_2_ dilatory action on the cerebral vasculature remains unclear^9^, conventional pharmacological or genetic approaches cannot be applied with certainty. Therefore, to study the role of CO_2_ in the development of the neurovascular response, we used an alternative experimental strategy which involved saturation of the brain CO_2_-sensitive mechanism(s) with exogenous CO_2_ given in the inspired air – a condition referred herein as hypercapnia (Figure 1a). We reasoned that if CO_2_ acts as a signalling molecule between brain neurons and the vasculature then in conditions of surplus exogenous CO_2_ any extra metabolic CO_2_ generated as a result of increased neuronal activity should have no further effect on the cerebral vasculature and the neurovascular response should be blocked (Figure 1a). However, if mechanisms other than CO_2_-mediated signalling mediate the neurovascular response, then its development should not at all be affected by hypercapnia.

**Figure 1.**
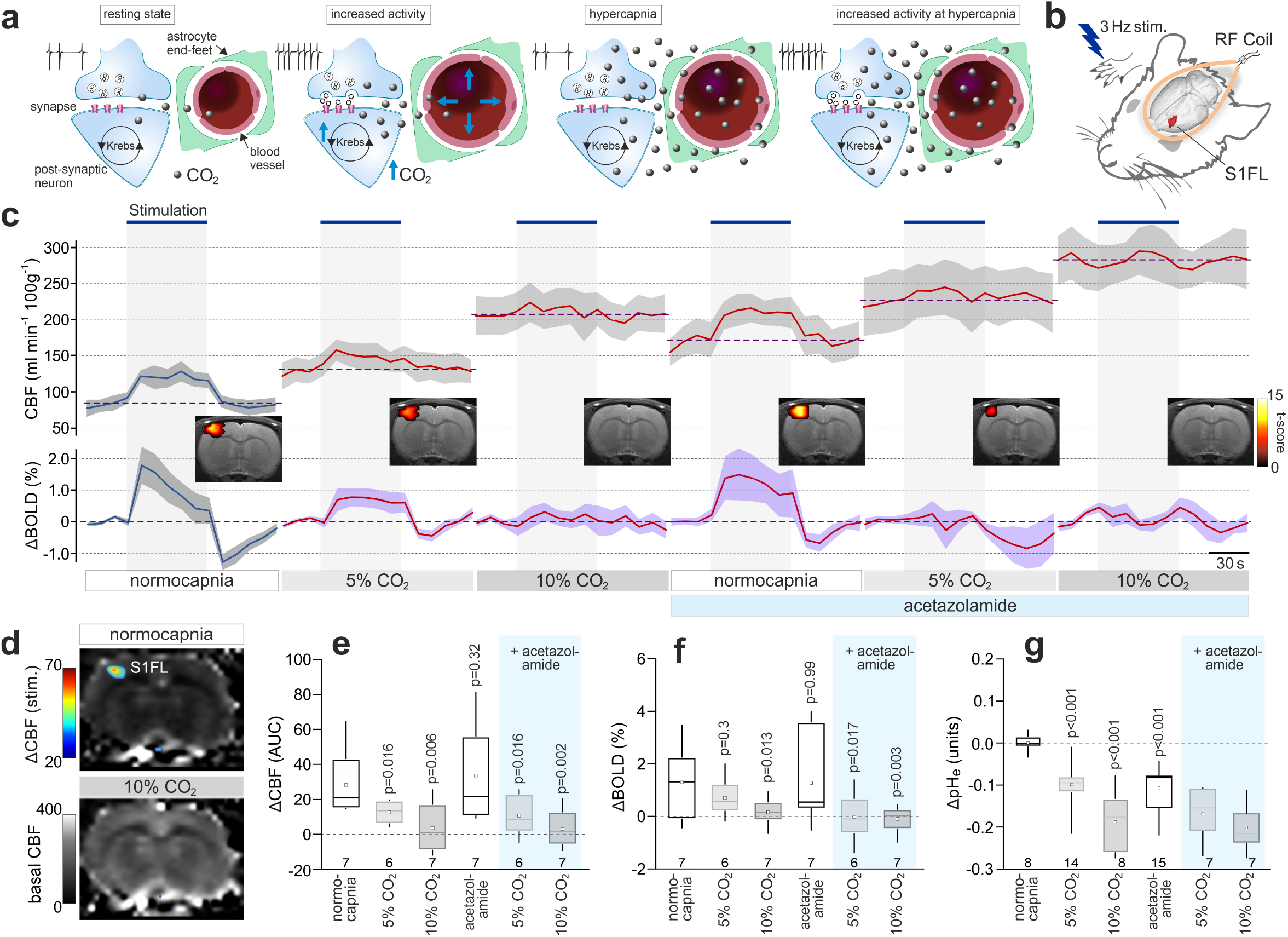
Exogenous CO_2_ prevents the development of the neurovascular response in the somatosensory cortex in rats. **a,** Schematic depiction of the neurovascular unit illustrating the central hypothesis and the experimental approach taken to study the potential role of CO_2_ as a signalling molecule of the neurovascular coupling response. Surplus of exogenous CO_2_ was given in the inspired air (hypercapnia) to saturate the intrinsic CO_2_-sensitive mechanism. If the hypothesis is correct then in the presence of surplus of exogenous CO_2_, any extra metabolic CO_2_ generated as a result of increased neuronal activity should have no additional effect on local blood flow and the neurovascular response should be blocked. If the mechanisms other than CO_2_ signalling mediate the neurovascular response, then the response should not be affected by exogenous CO_2_. **b,** Cerebral blood flow (CBF) and blood oxygen level dependent (BOLD) signals in the forepaw region of the somatosensory cortex (S1FL) in anesthetized rats were recorded using an arterial spin labelling (ASL) sequence with T2* weighted imaging. Somatosensory pathways were activated by electrical stimulation of the forepaw. **c,** CBF and BOLD responses in the S1FL region induced by electrical forepaw stimulation (3 Hz, 1.5 mA) at baseline, in conditions of 5% and 10% inspired CO_2_, after the administration of carbonic anhydrase inhibitor acetazolamide (10 mg kg^-1^, i.v.) and in conditions of 5% and 10% inspired CO_2_, applied concomitantly with systemic carbonic anhydrase inhibition with acetazolamide. *Insets* show representative activation maps illustrating mean BOLD signal changes in response to activation of somatosensory pathways. Colour bar: *t*-score from statistical paramagnetic mapping mixed-effects analysis, *p*<0.05 (uncorrected). **d,** Representative ASL images illustrating CBF at rest (normocapnia) and in conditions of 10% inspired CO_2_. Overlaid (false colour scale) illustrates CBF response to forepaw stimulation. **e,f,** Summary data illustrating the effect of 5% and 10% inspired CO_2_ given before and after systemic administration of acetazolamide on CBF and BOLD responses recoded in the S1FL region of the cortex. **g,** Summary data illustrating the effect of acetazolamide, as well as 5% and 10% inspired CO_2_ given before and after systemic administration of acetazolamide, on extracellular pH recorded using fast scan cyclic voltammetry in the S1FL region. *P* values, ANOVA.

In this study we used laboratory rats and implemented an arterial spin labelling (ASL) fMRI sequence with T2* weighted imaging for combined measurement of local CBF (in ml 100 g^-1^ min^-1^) and BOLD signal changes in the forelimb region 1 of the somatosensory cortex (S1FL) (Figure 1b). Under normocapnic conditions (PaCO_2_ ~35 mmHg) activation of somatosensory pathways by electrical forepaw stimulation triggered robust CBF increases (from 88±10 to 120±11 ml 100 g^-1^ min^-1^, p=0.015, n=7) and BOLD responses (1.3±0.5%, p<0.05, n = 7) in the S1FL region (Figure 1c). The addition of 5% and 10% inspired CO_2_ increased global CBF (Figure 1c,d) and reduced the amplitude of the CBF responses to somatosensory stimulation by 55% (p=0.016) and 87% (p=0.006), respectively (Figure 1c,e). The BOLD response was reduced by 83% (0.33±0.2%, p=0.013) in conditions of 10% inspired CO_2_ (Figure 1c,f).

Next, we determined whether the loss of the neurovascular response during hypercapnia is due to acidification of the brain tissue derived from CO_2_ hydration. We manipulated the brain extracellular pH by systemic administration of a carbonic anhydrase inhibitor acetazolamide (10 mg kg^-1^, i.v.), which is well known to reduce the brain pH. Acetazolamide-induced extracellular acidification in the S1FL region was similar in magnitude to acidification induced by 5% inspired CO_2_ (decrease by 0.10±0.02 pH units after acetazolamide *vs* −0.11±0.01 pH units in conditions of 5% inspired CO_2_; Figure 1g). Acetazolamide increased the basal CBF but had no effect on the magnitude of the CBF and BOLD responses induced by somatosensory stimulation (Figure 1c,e,f and Supplementary Figure 1). This observation is consistent with the data obtained in humans^10^. Acetazolamide also had no effect on CO_2_-induced increases in CBF (Figure 1g and Supplementary Figure 2). Yet, CO_2_ given in the inspired air still effectively blocked the neurovascular response in conditions of systemic acetazolamide action (Figure 1c,e,f).

Analysis of the evoked neuronal activity in the S1FL region revealed the inhibitory effect of CO_2_. The neuronal responses were reduced by 34% (p=0.04) and 48% (p=0.001) in conditions of 5% and 10% inspired CO_2_ (Figure 2f and Supplementary Figure 3). Interestingly, acetazolamide had no effect (p=0.99) on the evoked neuronal activity (Supplementary Figure 3), suggesting that the inhibitory effect of CO_2_ is independent of its effect on brain pH. Increasing the intensity of forepaw stimulation two-fold from 1.5 mA to 3 mA fully compensated (and exceeded the responses evoked by 1.5 mA stimulation) for the CO_2_-induced inhibition (Figure 2f and Supplementary Figure 3), yet failed to restore the neurovascular response, with both the CBF and BOLD responses remained markedly suppressed by 10% inspired CO_2_ (by 54%, p=0.020 and 61%, p=0.018, respectively; Figure 2c-e).

**Figure 2.**
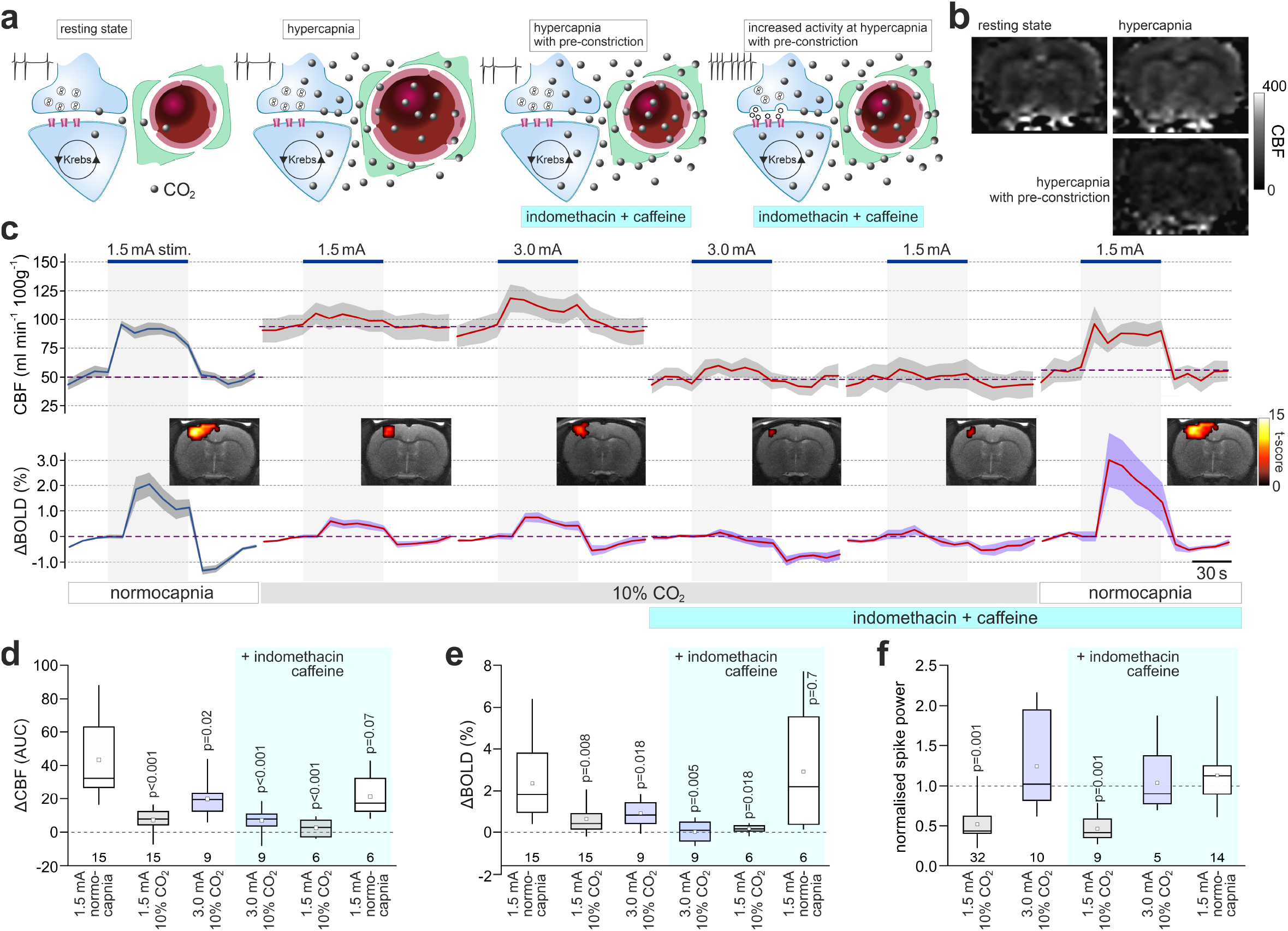
Exogenous CO_2_ prevents the development of the neurovascular response in the somatosensory cortex independently of basal CBF. **a,** Schematic depiction of the neurovascular unit illustrating the experimental approach used to control for the effect of exogenous CO_2_ on basal CBF. Extra CO_2_ generated during enhanced neuronal activity has no effect on local blood flow in the presence of surplus exogenous CO_2_. However, under these conditions the vasculature is dilated and may lack the capacity to respond further. To counteract the dilatory action of 10% inspired CO_2_ on brain vasculature, the animals were given a combination of caffeine and indomethacin (both at 10 mg kg^-1^, i.v.) to reduce the CBF to the baseline level recorded at normocapnia. **b,** Representative ASL images illustrating CBF at rest (normocapnia), in conditions of 10% inspired CO_2_, and after the systemic treatment with caffeine/indomethacin during hypercapnia. **c,** CBF and BOLD responses in the S1FL region induced by forepaw stimulation at baseline, in conditions of 10% inspired CO_2_, after the systemic administration of caffeine and indomethacin in conditions of 10% inspired CO_2_ and after withdrawal of inspired CO_2_. To compensate for CO_2_-induced inhibition of the neuronal activity, the intensity of forepaw stimulation was increased two-fold from 1.5 mA to 3 mA when 10% inspired CO_2_ was applied. *Insets* show representative activation maps illustrating mean BOLD signal changes in response to forepaw stimulation. Colour bar: *t*-score from SPM mixed-effects analysis, *p*<0.05 (uncorrected). **d,e,** Summary data illustrating CBF and BOLD responses in the S1FL region induced by electrical forepaw stimulation at baseline, in conditions of 10% inspired CO_2_, at 3 mA stimulus intensity in conditions of 10% inspired CO_2_, after the systemic administration of caffeine/indomethacin in conditions of 10% inspired CO_2_ and after the withdrawal of inspired CO_2_. **f,** Summary data illustrating the effect of each of the experimental conditions on the evoked neuronal responses (expressed as spike power) in the S1FL region triggered by electrical forepaw stimulation. Integrated spike activity is normalized and presented relative to the responses recorded at baseline. *P* values, ANOVA.

As expected, experimental hypercapnia increased the global CBF (Figure 1c,d, 2b,c; Supplementary Figure 4). Therefore, it could be argued that in these conditions neuronal activity fails to trigger a neurovascular response over and above the elevated CBF because the cerebrovascular reserve is exhausted by the potent dilatory action of CO_2_ on brain vasculature. To counteract this action of CO_2_ and lower the CBF to the baseline level recorded in normocapnia, we next gave the animals 10% CO_2_ in the inspired air and applied pharmacological agents known to reduce the CBF – caffeine^11^ and indomethacin^12^ (Figure 2a-c). The actions of neither caffeine (10 mg kg^-1^, i.v.) nor indomethacin (10 mg kg^-1^, i.v.) given alone were able to fully offset the effect of CO_2_ and reduce the CBF to the baseline level (Supplementary Figure 5). However, combined treatment with caffeine and indomethacin during hypercapnia effectively restored the baseline (normocapnic) level of CBF (49±6 ml 100 g^-1^ min^-1^ in 10% inspired CO_2_ following caffeine/indomethacin administration *vs* 53±4 ml 100 g^-1^ min^-1^ at baseline; p=0.77) (Figure 2b,c). Although in conditions of 10% inspired CO_2_ the baseline CBF was fully restored by this treatment, no significant CBF and BOLD responses to somatosensory stimulation were recorded in the S1FL region (reduction by 84%, p<0.001 and 98%, p=0.005; respectively; Figure 2c-e). In the continuing presence of systemic caffeine and indomethacin, the CBF and BOLD responses to somatosensory stimulation rapidly and fully recovered after the withdrawal of inspired CO_2_, indicating that the actions of these drugs had no effect on the neurovascular response (Figure 2c-e).

All penetrating and intraparenchymal cerebral blood vessels are wrapped by end-feet of astrocytes (Figure 3a). In response to increases in the neuronal activity astrocytes release bicarbonate anion (HCO_3_^-^) via electrogenic sodium-bicarbonate cotransporter 1 (NBCe1) and, by doing so, help to maintain local brain acid-base balance^13^. In mice with astrocyte-specific genetic deletion of NBCe1, brain CO_2_/pH homeostasis is disrupted^13^. We next hypothesized that if CO_2_ mediates the neurovascular coupling response than this response should be affected in conditions when CO_2_/pH homeostasis is not effectively maintained (Figure 3a). Using the minimally invasive method of fast cyclic voltammetry^14^ to measure changes in brain tissue PO_2_ (Figure 3b), we recorded stereotypical neurovascular response in the S1FL region evoked by somatosensory stimulation in mice. In control animals, increases in brain tissue PO_2_ (peak 4.6±1.5 mmHg; n=7; p=0.018) were recorded 25 s after the onset of stimulation, returning to the baseline within ~1 min after the end of the stimulation (Figure 3d). In conditions of NBCe1 knockdown in astrocytes (Figure 3c), the neurovascular response in the S1FL region induced by forepaw stimulation was completely abolished, with tissue PO_2_ falling below the baseline during the period of stimulation (by −3.5±1.8 mmHg; n=7; p=0.010), resulting in a marked difference in the integral PO_2_ response between the experimental groups (−7.5±5.0 mmHg.s in NBCe1 knockout *vs* 15.8±5.0 mmHg.s in the control animals; p=0.007) (Figure 3d,e).

**Figure 3.**
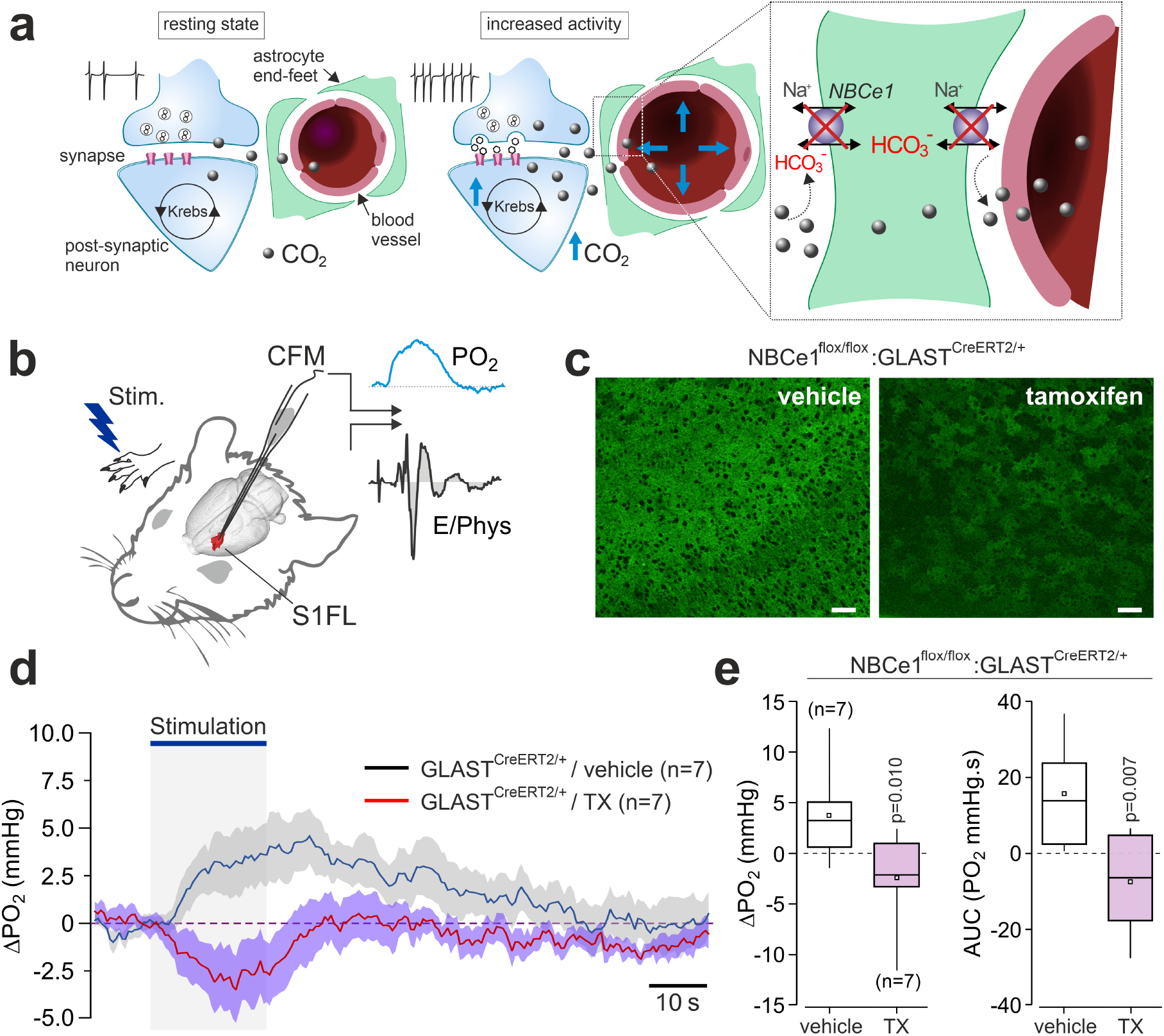
Genetic knockdown of sodium bicarbonate cotransporter 1 (NBCe1) expression in astrocytes blocks the neurovascular response in the somatosensory cortex. **a,** Schematic depiction of the neurovascular unit illustrating disrupted CO_2_/HCO_3_-transport in astroglial endfeet in conditions of NBCe1 deficiency. **b**, Changes in tissue PO_2_ and the neuronal activity in the S1FL cortical region of mice were recorded using fast cyclic voltammetry. The carbon fibre electrode was placed in the S1FL region. The forepaw was stimulated electrically to activate the somatosensory pathways. Switching between current and voltage recording allowed near-simultaneous detection of brain tissue PO_2_ and the evoked neuronal activity. **c,** Knockdown of NBCe1 expression in cortical astrocytes. Confocal images illustrate immunohistochemical detection of NBCe1 in the cortex of NBCe1^flox/flox^:GLAST^CreERT2/+^ mice treated with the vehicle (oil) or tamoxifen. Tamoxifen treatment of NBCe1^flox/flox^:GLAST^CreERT2/+^ mice resulted in a mosaic pattern of NBCe1 expression. Scale bars = 40 μm. **d**, Time course of tissue PO_2_ changes in the S1FL region induced by activation of somatosensory pathways (electrical stimulation of the contralateral paw; 3 Hz, 1.5 mA) in NBCe1^flox/flox^:GLAST^CreERT2/+^ mice treated with the vehicle or tamoxifen. NBCe1 knockdown in astrocytes prevented the development of the neurovascular response. **e**, Summary data illustrating peak and integral changes in PO_2_ evoked in the S1FL region by electrical forepaw stimulation in NBCe1^flox/flox^:GLAST^CreERT2/+^ mice treated with the vehicle or tamoxifen. *P* values – Mann Whitney-U test.

These results support the hypothesis that CO_2_ functions as one of the key signalling molecules responsible for the neuronal activity-dependent increases in local brain blood flow. The data obtained show that saturation of the brain CO_2_-sensitive vasodilatory mechanism by the provision of exogenous CO_2_, or inhibition of HCO_3_^-^ transport in astrocytes, leading to a disruption of brain CO_2_/pH homeostasis, block the development of the neurovascular response, independently of baseline CBF and the effects of CO_2_ on brain tissue pH and neuronal activity. This hypothesis is not incompatible with the current view of the neuronal activity-dependent control of brain arterioles and capillaries by arterial smooth muscle cells and pericytes^5^. In fact, the responses of the latter are mediated predominantly by the actions of ATP^5^, well known to be released by astrocytes and possibly other brain cells in a CO_2_-sensitive manner^15–18^. Yet, based on the results obtained in this study we suggest a reconsideration of the prevailing view^19^ that the neurovascular response is driven exclusively by neurotransmitters and/or K^+^ released by active neurons. We provide the first experimental data for a revised model which combines neurotransmitter-mediated ‘feed-forward’ and CO_2_-mediated metabolic feed-back mechanisms of the neurovascular coupling response working in concert for the purpose of efficient removal of CO_2_ generated by the brain.

## Methods

All animal experiments were performed in accordance with the European Commission Directive 2010/63/EU (European Convention for the Protection of Vertebrate Animals used for Experimental and Other Scientific Purposes) and the UK Home Office (Scientific Procedures) Act (1986) with project approval from the respective Institutional Animal Care and Use Committees. The animals were group-housed and maintained on a 12-h light cycle (lights on 07:00) and had *ad libitum* access to water and food.

### General animal preparation

Rats (Sprague-Dawley, 280-350 g) were anesthetized with α-chloralose (induction: 100 mg kg^-1^, maintenance: 30 mg kg^-1^ hr^-1^, i.v.). The femoral artery and vein were cannulated for continuous monitoring of the arterial blood pressure and the administration of anesthetic, respectively. Adequate depth of anesthesia was confirmed by the stability of arterial blood pressure and heart rate recordings, which did not show responses to a paw pinch. The animal was intubated and mechanically ventilated with oxygen-supplemented air using a small rodent ventilator (tidal volume ~0.8 ml per 100 g of body weight; ~60 strokes min^-1^). Neuromuscular blockade was established following administration of gallamine (induction: 5 mg kg^-1^, maintenance 2 mg kg^-1^ h^-1^, i.v.). Arterial PO_2_, PCO_2_, and pH were measured regularly and kept within the physiological ranges (PO_2_ 100-120 mmHg; PCO_2_ 35-40 mmHg; and pH 7.35-7.45) by adjusting the tidal volume and/or ventilator frequency as well as the amount of supplemental oxygen. Body temperature was maintained at 37.0 ± 0.5 °C using a servo-controlled heating pad.

### Functional magnetic resonance imaging

fMRI was used to record the neurovascular responses in the somatosensory cortex. We implemented an ASL sequence with T2* weighted imaging for combined measurement of local CBF (in ml 100 g^-1^ min^-1^) and BOLD signal changes in the S1FL region of the cortex. The measurements of absolute changes in CBF were essential to address the objectives of this study as the BOLD signal reflects an unknown combination of changes in local CBF, blood volume and CMRO_2_. Arterial spin labelling (ASL) measurements provided absolute values of cerebral perfusion to control for possible confounding effects of the experimental manipulations on the expression of the neurovascular response.

fMRI was performed using a 9.4 T Agilent horizontal bore scanner (Agilent) as described in detail previously^20,21^. The animal was anaesthetized and instrumented as described above *(General animal preparation)* and then transferred to the MRI scanner bed. The head was secured with ear and incisor bars. A 72 mm inner diameter volume coil was used for radio frequency transmission and signal was received using a 4-channel array head coil (Rapid Biomedical). First, a high-resolution anatomical reference scan was acquired using a fast spin echo sequence (TR/TE_eff_ = 3100/48 ms, ETL = 8, matrix size = 256 × 256, FOV = 35 mm × 35 mm, 30 slices, 1 mm slice thickness). The anatomical reference image was used to manually position the single functional coronal imaging slice (see below) to be centred on the S1FL region. Resting CBF and CBF/BOLD responses to somatosensory stimulation were recorded using a flow-sensitive alternating inversion recovery (FAIR) ASL sequence, with concurrent BOLD T2* weighted BOLD weighting, using the following sequence parameters: single shot gradient echo EPI readout, TR = 5000 ms, inflow time (TI) = 2000 ms, matrix size = 64 × 64 voxels, FOV = 35 × 35 mm, TE = 10 ms, single slice (slice thickness = 2 mm), inversion pulse bandwidth = 20,000 Hz. Maps of CBF were generated by fitting the data to previously established models^22^, where T1 in the cortex was assumed to be 1.7 s, the arterial transit time to be 0.3 s and the temporal duration of the tagged bolus to be 2 s^23^, based on previous measurements using the identical experimental conditions and equipment.

### fMRI data analysis

CBF and BOLD time-series data were extracted from manually drawn regions of interest (ROIs) based on the baseline functional data and fixed for all the subsequent conditions. The BOLD signal was taken from the alternate ‘control’ images in the FAIR ASL acquisition. In order to partially correct the ASL (ΔM) signal for marked BOLD T2* weighted signal changes that occur when the forepaw stimulus begins/ceases at a time between the labelled and control acquisitions^24^, the labelled image directly after the stimulus begins was substituted by the previous labelled image (captured during the baseline period). Similarly, the control image taken directly after the cessation of the stimulus was substituted by the previous control image (captured during the stimulation period). For the CBF and BOLD time-course data, occasional spike artefacts in the recorded signal due to hardware instability were removed using an automated de-spiking algorithm. BOLD signal responses are expressed as changes from the average of pre-stimulus baseline. CBF responses were assessed by calculating the area under the curve for each experimental condition. In order to generate maps of BOLD signal responses for visualisation purposes, the control images were spatially smoothed (0.5 mm FWHM Gaussian kernel), and first level analysis of each time-series using an on/off regressor derived from the applied forepaw stimulus paradigm (and convolved with the standard HRF (SPM)) was applied to generate the statistical maps. In order to visualize the BOLD activation maps to forepaw stimulation, a minimum threshold of p<0.0001 with a cluster size of >5 voxels was applied.

### Recordings of extracellular pH and neuronal activity

Changes in extracellular pH and evoked neuronal activity were recorded using fast scan cyclic voltammetry^14^ in a separate cohort of rats (n=27) under the experimental conditions that were identical to that during the imaging experiments *(General animal preparation).* Extracellular pH changes were recorded continuously during each of the experimental challenges (provision of CO_2_ in the inspired air and/or administration of pharmacological agents) and expressed as averages of values recorded during a 1 min period immediately prior to forepaw stimulation. Changes in the evoked neuronal activity were assessed by integration of the evoked volley of extracellular potentials with the baseline noise subtracted. Recordings obtained during three stimulus trains were averaged for the analysis.

### Conditional NBCe1 knockdown in astrocytes

To induce conditional NBCe1 knockdown in astrocytes, mice carrying a loxP-flanked NBCe1 allele (NBCe1^flox/flox^)^25^ were crossed with the mice expressing an inducible form of Cre (Cre^ERT2^) under the control of astrocyte-specific GLAST promoter^26^. Tamoxifen (100 mg kg^-1^) dissolved in corn oil was given to NBCe1^flox/flox^:GLAST^CreERT2/+^ mice at postnatal week 7 and the expression level of NBCe1 was examined 6 weeks after the tamoxifen treatment. Breeding was organized trough PCR genotyping obtained from tail DNA biopsies. Astroglial recombination specificity of GLAST-CreERT2 mouse line has been reported in several prior studies^26,27^. NBCe1 expression in cortical astrocytes was reduced by ~40%^13^. Littermate NBCe1^flox/flox^:GLAST^CreERT2/+^ mice injected with tamoxifen vehicle (oil) were used as controls.

### Experimental protocols

*<u>Experiment 1</u>: The effect of exogenous CO_2_ on neuronal activity-dependent increases in local cerebral blood flow in the somatosensory cortex.* In rats, anaesthetized and instrumented as described above *(General animal preparation),* activation of somatosensory pathways leading to robust CBF and BOLD signal responses in the S1FL cortical region was achieved by electrical stimulation of the forepaw (300 μs pulse width, 3 Hz, 1.5 mA), applied using bipolar subcutaneous electrodes delivering fixed-current pulses from an isolated stimulator (Digitimer DS3). Three repeated forepaw stimulations 60 s in duration, with a 60 s inter-stimulus interval, and 60 s baseline period were applied, with CBF and BOLD signal responses to stimulations averaged and compared between each of the following experimental conditions: 1) <u>control condition</u>: animals were ventilated with oxygen-enriched room air and ventilation parameters adjusted to ensure the arterial PO_2_, PCO_2_ and pH were maintained within the physiological ranges, as described above; 2) <u>5% inspired CO_2_ condition</u>: 5% CO_2_ was added to the inspired gas mixture; 3) <u>10% inspired CO_2_ condition</u>: 10% CO_2_ was added to the inspired gas mixture; 4) <u>brain acidosis condition</u>: inspired CO_2_ was withdrawn and acetazolamide was given systemically (10 mg kg^-1^, i.v.) to inhibit carbonic anhydrase (CA) causing a decrease in brain pH; 5) <u>5% inspired CO_2_ in conditions of CA inhibition</u>: in the continuing presence of systemic acetazolamide, 5% CO_2_ was added to the inspired gas mixture; 6) <u>10% inspired CO_2_ in conditions of CA inhibition</u>: in the continuing presence of systemic acetazolamide, 10% CO_2_ was added to the inspired gas mixture. Each condition was established for at least 10 min prior to the electrical stimulation of the forepaw to trigger the neurovascular response in the S1FL region.

*<u>Experiment 2</u>: The effect of exogenous CO_2_ on neuronal activity-dependent increases in local cerebral blood flow in the somatosensory cortex before and after the baseline (normocapnic) level of CBF is restored pharmacologically.* To counteract the vasodilatory effect of CO_2_ and lower the CBF to the baseline level recorded at normocapnia, we next gave the animals 10% CO_2_ in the inspired air and then applied two pharmacological agents known to reduce the CBF, – caffeine and indomethacin. These drugs were applied either alone or in combination, in 3 separate experiments involving a sequence of experimental conditions described below. Three repeated forepaw stimulations 60 s in duration, with a 60 s inter-stimulus interval, and 60 s baseline period were applied with CBF and BOLD signal responses to stimulations averaged and compared between each of the following experimental conditions, studied sequentially: 1) <u>control condition</u>: animals were ventilated with oxygen-enriched room air and ventilation parameters adjusted to ensure the arterial PO_2_, PCO_2_ and pH were maintained within the physiological ranges, as described above; 2) <u>10% inspired CO_2_ condition</u>: 10% CO_2_ was added to the inspired gas mixture; 3) <u>10% inspired CO_2_ condition with baseline (normocapnic) CBF restored pharmacologically</u>: either caffeine, indomethacin or a combination of caffeine and indomethacin (both drugs at 10 mg kg^-1^, i.v.) were given concurrently with 10% CO_2_ in the inspired gas mixture; 4) <u>systemic caffeine/indomethacin condition</u>: inspired CO_2_ was withdrawn in conditions of continuing systemic caffeine/indomethacin action.

A separate group of animals was studied under the same sequence of experimental conditions but when 10% CO_2_ was given in the inspired gas mixture (conditions 2 and 3), the electrical current applied to the forepaw was increased to 3 mA in order to compensate for the CO_2_-induced inhibition of the neuronal activity, as described in the main text.

*<u>Experiment 3</u>: The effect of astrocyte-specific NBCe1 knockdown on neurovascular coupling response in the somatosensory cortex.* Fast scan cyclic voltammetry was used to record the neurovascular response in the S1FL region of the somatosensory cortex in mice. The expression of the neurovascular response was assessed from the recordings of changes in brain tissue PO_2_, previously reported to accurately match BOLD responses to somatosensory stimuli^14^. Mice (3-4 mo old of both sexes) were anesthetized with a mixture of ketamine/xylazine (100 mg kg^-1^/10 mg kg^-1^, i.p.). The head was secured in a stereotaxic frame with ear and incisor bars. Oxygen-enriched room air was supplied to the animal breathing unaided throughout the experiment. After a small craniotomy (~1 mm^2^), carbon fibre microelectrode (∅ 7 μm) was advanced into the S1FL cortex until evoked field potentials were recorded in response to the electrical stimulation (1 Hz) of the contralateral forepaw. A series of voltage ramps (200 V s^-1^) from 0 to −1 V were applied to the electrode at a frequency of 2 Hz. The recording was amplified, digitised and stored for offline isolation of faradaic current to determine changes in brain tissue PO_2_.

Electrical forepaw stimulations (300 μs pulse width, 3 Hz, 1.5 mA) were applied to activate somatosensory pathways and evoke the neurovascular response in the S1FL region. Cortical tissue PO_2_ responses evoked during the 3 stimulations (each 20 s long) were averaged within each animal to obtain a profile of the response for each subject. Peak of the response (difference in PO_2_ between the response maximum and the pre-stimulus baseline, expressed in mmHg) and integral (area under the curve, expressed in mmHg.s) were compared between the astrocyte specific NBCe1 knockout (NBCe1^flox/flox^:GLAST^CreERT2/+^ animals given tamoxifen) and control (NBCe1^flox/flox^:GLAST^CreERT2/+^ animals given vehicle) mice.

### Drugs

Acetazolamide and indomethacin (Cambridge Bioscience) were dissolved in 100% DMSO (100 mg ml^-1^) and administered i.v. (10 mg kg^-1^). Caffeine (Sigma) was dissolved in saline and administered i.v. (10 mg kg^-1^). The maximum volume of administered DMSO was 100 μl kg^-1^, previously shown to have no effect on cerebrovascular responses^28^.

### Data analysis

The data are shown as means ± s.e.m. or as box and whisker plots. Normal distribution of data was determined by the Shapiro-Wilk test. Grouped data were analyzed using ANOVA or Friedman Test (for non-normally distributed data) when comparing data between more than two groups. Post-hoc analysis was performed corrected for multiple comparisons using Tukey’s or Dunn’s method. Wilcoxon’s signed rank tests was used to compare the data between two experimental groups.

## Acknowledgements

This work was supported by The Wellcome Trust (A.V.G.) and the Fondecyt Iniciación Grant 11190678 (I.R.). A.V.G is a Wellcome Trust Senior Research Fellow (Ref: 200893). CECs is funded by the Chilean Government through the Centers of Excellence Base Financing Program.

