## Supplementary Figures 1-5 for "CO_2_ signalling mediates neurovascular coupling in the cerebral cortex"

Hosford et al.

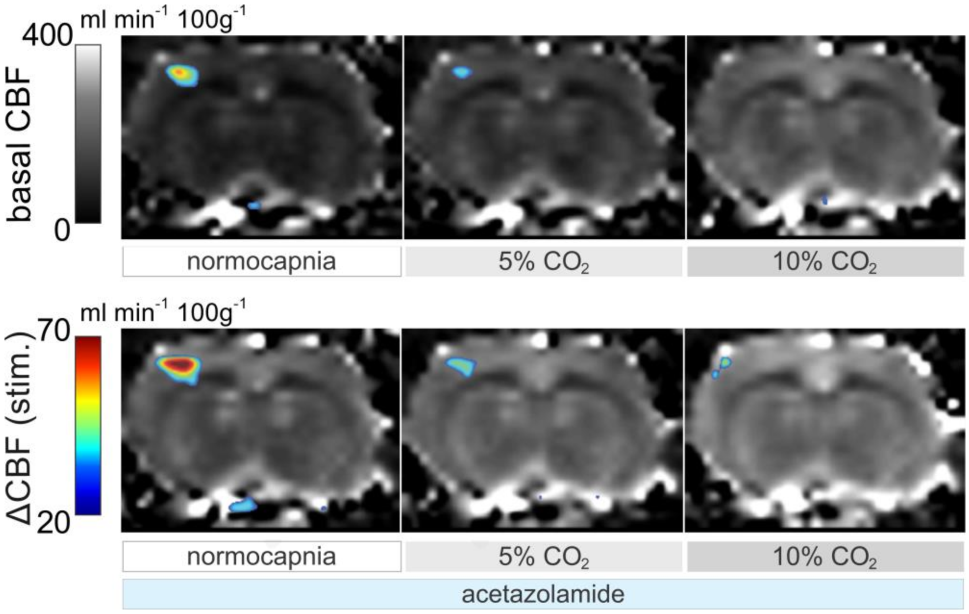


Supplementary Figure 1| Representative arterial spin labelling images of the rat brain illustrating cerebral blood flow (CBF) at baseline (normocapnia), in conditions of 5% and 10% inspired $CO_2$, after the administration of carbonic anhydrase inhibitor acetazolamide (10 mg $kg^{-1}$, i.v.) and in conditions of 5% and 10% inspired $CO_2$, applied concomitantly with systemic carbonic anhydrase inhibition with acetazolamide. Overlaid (false colour scale) illustrates CBF response in the S1FL region of the somatosensory cortex induced by electrical forepaw stimulation (3 Hz, 1.5 mA).

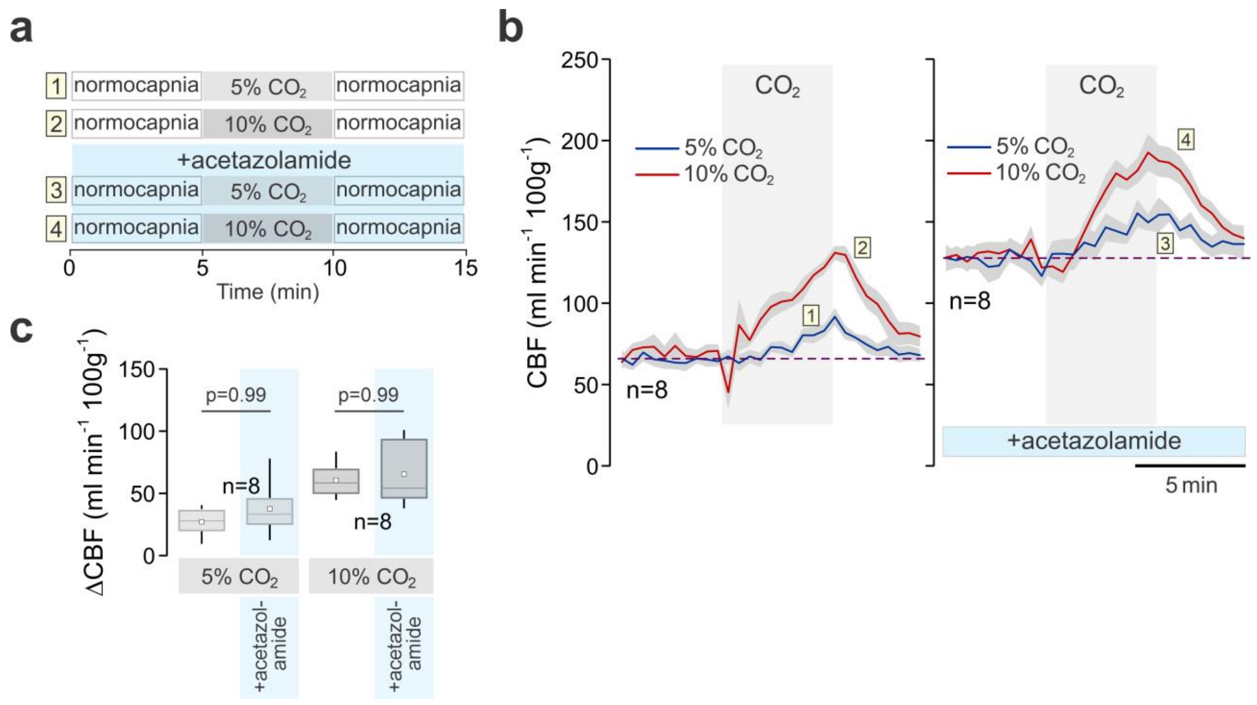


Supplementary Figure 2| Cerebrovascular reactivity to $CO_2$ is not affected by systemic carbonic anhydrase inhibition with acetazolamide. **a**, Schematic depiction of the experimental timeline. $CO_2$ challenge (5% or 10% $CO_2$ in the inspired air) was given for 5 min before and after systemic administration of acetazolamide (10 mg $kg^{-1}$). **b**, Time-course of the whole brain CBF changes recorded using arterial spin labelling MRI in anaesthetised rats at resting conditions and in response to 5% and 10% inspired $CO_2$, applied before and after systemic carbonic anhydrase inhibition with acetazolamide. Numbers refer to the experimental conditions depicted at the schematic shown in **a**; **c**, Summary data illustrating peak absolute $CO_2$-induced increases in CBF from the baseline before and after administration of acetazolamide. *P* values, Mann Whitney-U test.

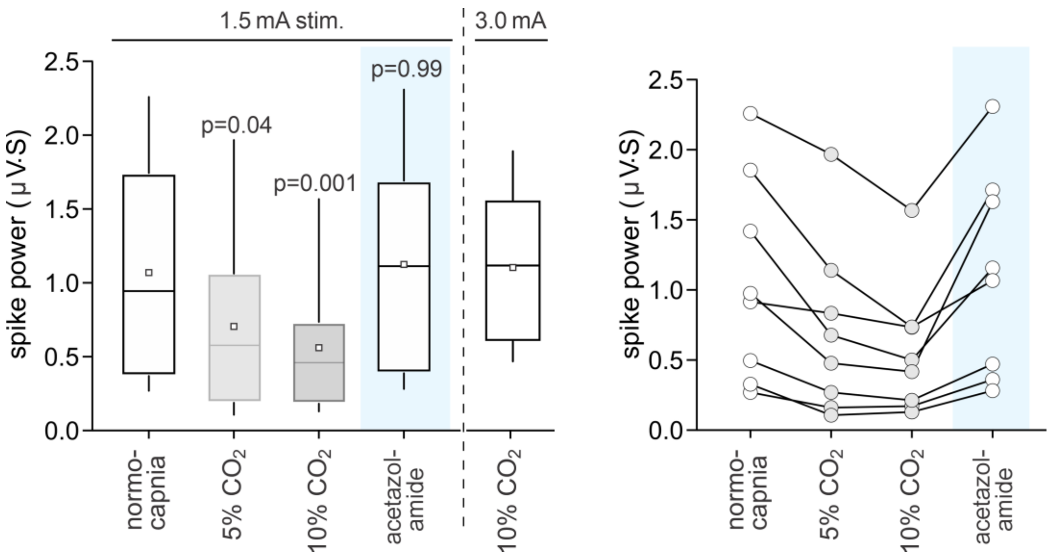


Supplementary Figure 3| $CO_2$ reduces neuronal excitability in the somatosensory cortex. Summary data (box and whisker plots and individual data) illustrating the effect of 5% and 10% inspired $CO_2$ and systemic carbonic anhydrase inhibition with acetazolamide (10 mg $kg^{-1}$) on the evoked neuronal responses (expressed as spike power) in the S1FL region of the somatosensory cortex induced by electrical forepaw stimulation. Acetazolamide had no effect on the neuronal activity, suggesting that the inhibitory effect of $CO_2$ is independent of its effect on brain pH. *P* values, ANOVA.

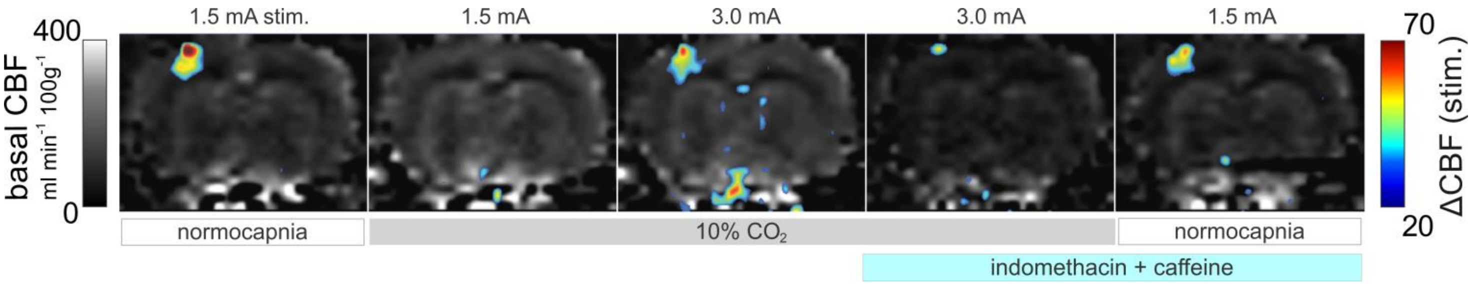


Supplementary Figure 4| Representative arterial spin labelling images of the rat brain illustrating CBF at baseline (normocapnia), in conditions of 10% inspired $CO_2$, after the systemic administration of indomethacin and caffeine (both at 10 mg $kg^{-1}$; i.v.) in conditions of 10% inspired $CO_2$, and after the withdrawal of inspired $CO_2$. Overlaid (false colour scale) illustrates CBF responses in the S1FL region of the somatosensory cortex induced by electrical forepaw stimulation (3 Hz, 1.5 or 3 mA).

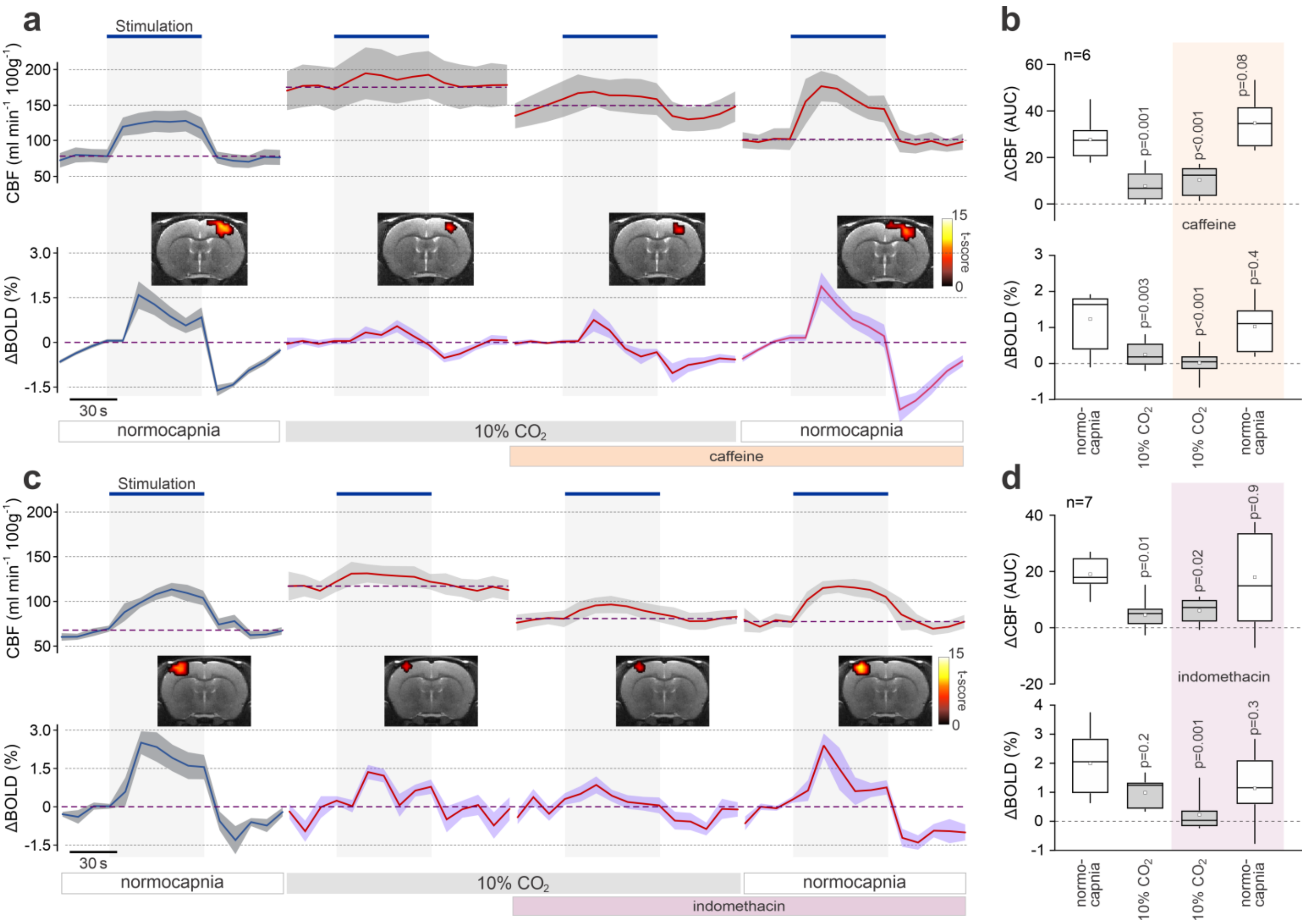


Supplementary Figure 5| Exogenous $CO_2$ prevents the development of the neurovascular response in the somatosensory cortex independently of basal CBF. **a**, CBF and BOLD responses in the S1FL region induced by electrical forepaw stimulation at baseline, in conditions of 10% inspired $CO_2$, after the systemic administration of caffeine (10 mg $kg^{-1}$; i.v.) in conditions of 10% inspired $CO_2$ and after withdrawal of inspired $CO_2$. **b,** Summary data illustrating integral CBF and peak BOLD responses in the S1FL region induced by electrical forepaw stimulation at baseline, in conditions of 10% inspired $CO_2$, after the systemic administration of caffeine in conditions of 10% inspired $CO_2$ and after the withdrawal of inspired $CO_2$. **c**, CBF and BOLD responses in the S1FL region induced by electrical forepaw stimulation at baseline, in conditions of 10% inspired $CO_2$, after the systemic administration of indomethacin (10 mg $kg^{-1}$; i.v.) in conditions of 10% inspired $CO_2$ and after withdrawal of inspired $CO_2$. **d,** Summary data illustrating integral CBF and peak BOLD responses in the S1FL region induced by electrical forepaw stimulation at baseline, in conditions of 10% inspired $CO_2$, after the systemic administration of indomethacin in conditions of 10% inspired $CO_2$ and after the withdrawal of inspired $CO_2$. Activation maps illustrate mean BOLD signal changes in response to forepaw stimulation. Colour bars: *t*-score from SPM mixed-effects analysis, $p<0.05$ (uncorrected). *P* values, ANOVA.
